# Evaluation of the growth performance, physiological traits, antioxidant indices, and heat shock protein 70 to dietary supplementation of stinging nettle (*Urtica dioica*) in broilers exposed to chronic heat stress

**DOI:** 10.1101/2021.02.26.433121

**Authors:** Mehrad Mirsaiidi Farahani, Seyedeh Alemeh Hosseinian

## Abstract

Heat stress is known as one of the most prevalent environmental stressors in poultry production, which is associated with oxidative stress. Stinging nettle is a medicinal herb with strong antioxidant properties. The present study was conducted to evaluate the effects of dietary stinging nettle at two different levels (2% and 4%) on growth performance and oxidative stress indices of broilers exposed to chronic heat stress. On day 14, a total of 240 broiler chickens were randomly assigned to 6 treatment groups as follows: 1) NC: negative control; 2) HS: heat-stressed broiler; 3) HS-SN2: heat-stressed broiler fed by 2% dietary stinging nettle; 4) HS-SN4: heat-stressed broilers fed by 4% stinging nettle; 5) SN2: no heat-stressed broilers fed by 2% dietary stinging nettle; 6) SN4: no heat-stressed broilers fed by 4% dietary stinging nettle. Diet supplementation with stinging nettle was performed from days 14 to 35 and a chronic heat stress was induced from days 22 to 29. The growth performance and oxidative indices were evaluated on days 14, 21, 29, and 35. Rectal temperature and panting frequency were assessed on days 22, 25, and 29. As a result, weight gain and food intake decreased in the HS compared to the NC, and these parameters increased in the HS-SN2 and HS-SN4 compared to the HS (*P*<0.05). The HS, HS-SN2, and HS-SN4 groups had a significantly higher rectal temperature and panting frequency. The HS had the higher circulating MDA and HSP70, and lower T-AOC, SOD, and GSH-Px compared to the treatments. The HS-SN4 had a significantly lower MDA and HSP70, and higher T-AOC, GSH-Px, and SOD compared to the HS and HS-SN2. In conclusion, the addition of 4% stinging nettle powder to the broilers’ diet improved the oxidative status in heat-stressed birds. Furthermore, this herb could be utilized as a feed additive in poultry diet to improve bird’s health and defense mechanisms under stressful conditions.

## Introduction

Heat stress is one of the most common environmental stressors in poultry production. Broilers are very susceptible to heat stress due to lack of sweat glands, abundant feathers, and high metabolic rate (Xie et al., 2015). The heat stress adversely affects the birds’ physiology, health, and productive performance, causing endocrine impairment, metabolic disturbance, and immunity depression (Wang et al., 2018). In addition, heat stress leads to imbalance between redox status and oxidative stress in birds (Zheng et al., 2018). It also increases the reactive oxygen species (ROS) generation and inactivates the antioxidant defense systems in the body cells of birds, which lead to cell damages by proteins oxidation, nucleic acids damage, and lipids peroxidation (Habibi et al., 2014). Heat stress-induced oxidative stress increases the susceptibility of broilers to infectious and non-infectious diseases (Rahmani et al., 2017). Several methods have been used to evaluate the oxidative status in broilers subjected to heat stress. Measuring the serum and tissue levels of malondialdehyde (MDA), as a lipid peroxidation marker, is believed to be one of the most common biomarkers for assessing oxidative status in stressed broilers (Nawab et al., 2018). Evaluating the tissue levels of heat shock proteins (HSPs) is another prevalent index in broilers under heat stress (Xu et al., 2017). Cellular expression of HSPs increases in response to physical, chemical, and biological stresses to protect the broilers against cellular damages following oxidative stress (Tang et al., 2018). HSP70 is a major HSP that combines with unfolded proteins to repair them and maintain the normal physiological function of cells (Cedraz et al., 2017). Previous studies have shown that the HSP70 levels in hepatic and intestinal cells increased in heat-stressed broilers (Xu et al., 2017).

Various mechanisms are involved in the body of broilers to neutralize ROS and protect the cells against heat stress-induced oxidative stress, including enzymatic antioxidant systems, such as glutathione peroxidase (GPx) and superoxide dismutase (SOD), and non-enzymatic antioxidant systems, glutathione (GSH) for instance (Lu et al., 2007). It was reported that the activity of body antioxidant systems is not sufficient in birds exposed to chronic heat stress and dietary antioxidant supplementation could be beneficial to mitigate the adverse effects of heat stress-induced oxidative stress (Nawab et al., 2019).

Various researchers have employed different dietary antioxidants for attenuating harmful effects of heat stress on poultry (Toghyani et al., 2011; Nawab et al., 2019). In the recent years, the use of antioxidant-rich plants as feed additives in the broiler diet has also gained popularity among researchers owing to their various pharmacological and beneficial effects on health, oxidant status, and productive performance of broilers (Akbarian et al., 2016). One of these medicinal herbs is stinging nettle (*Urtica dioica*), which has been abundantly utilized as a phytogenic feed additive in poultry diet (Loetscher et al., 2013a, b).

*Urtica dioica L*. (*U. dioica*, Urticaceae) is an herbaceous and perennial plant often known as stinging nettle or nettle. It has been confirmed that stinging nettle has strong antioxidant, anti-inflammatory, antiviral, and anticarcinogenic properties owing to its high content of terpenoids, phenylpropanoids, flavonol glycosides, carotenoids, tocopherol, and ascorbic acid (Kregiel et al., 2018). Feeding with fresh leaves of stinging nettle in broilers is common in certain parts of Europe due to its unique pharmacological and nutritional properties (Loetscher et al., 2013b). Adding stinging nettle to daily diet of broilers has led to an improvement in their well-being and health status (Kregiel et al., 2018). Some studies have reported that dietary supplementation of 2-4% of stinging nettle had positive effects on growth performance and body’s antioxidant capacity in broilers (Sharma et al., 2018). In addition, this herb exerts strong antioxidant effects via scavenging the free radical and increasing the activities of antioxidant enzymes in living organisms under stressors (Joshi et al. 2014). Ahmadipour and Khajali (2019) used dietary supplementation of stinging nettle powder to attenuate oxidative stress induced by pulmonary hypertension syndrome.

According to previous findings, it was hypothesized that stinging nettle, owing to its antioxidant properties, may reduce the adverse effects of chronic heat stress in broilers; therefore, the present study was designed to evaluate the effects of dietary stinging nettle at two different levels (2% and 4%) on oxidative stress indices of broilers exposed to chronic heat stress. Furthermore, the effects of this plant on growth performance of heat stressed broilers were investigated. The results of the present study may also aid to better dietary management of under heat stress broilers.

## Materials and methods

### Birds and experimental groups

All the study protocols were approved by the Iranian Animal Ethics standards under the supervision of the Iranian Society for the Prevention of Cruelty to Animals and Shiraz University Research Council (IACUC no: 4687/63). A total of 240 one-day-old male *Arbor Acres* broiler chickens weighing 45±5 g were housed in stainless-steel cages (size of cage: 200 cm length ×100 cm width ×80 cm height) with 10 birds/m^2^ density, 6 nipple drinkers, and one feed trough (185 cm length) per cage in thermostatically controlled rooms with a relative humidity of about 60±10%. A continuous 24-h light exposure was performed for the first 3 days, and a 16L: 8D lighting schedule was applied from day 3 until end of the study. All the nutritional and environmental conditions were programmed based on *Arbor Acres* Broiler Management Guide (Aviagen, 2018).

The basal diet (Table 1) was formulated based on corn and soybean meal to meet the nutrient requirements of the broilers, according to National Research Council (NRC, 1994). All the broilers were fed *ad libitum* with a starter (days 0–10), grower (days 10–25), and finisher (days 25–35) diets. All the broilers were fed by a similar basal diet from day 1 to 14. On day 14, the birds were randomly divided into six equal groups (with 4 replicates of 10 birds in each group) and subjected to different feeding and thermal conditions as presented in Figure 1 and the following explanation: the negative control group (NC) was fed by basal diet and kept at ambient temperature (24±1°C); the heat-stressed group (HS) was fed by basal diet and reared under cyclic heat stress; the heat-stressed plus 2% and 4% stinging nettle groups (HS-SN2 and HS-SN4) were fed by basal diet supplemented with 2% and 4% stinging nettle powder, respectively, and were subjected to cyclic heat stress; 2% and 4% stinging nettle groups (SN2 and SN4) were fed by basal diet supplemented with 2% and 4% stinging nettle powder, respectively, without being exposed to heat stress.

**Table 1.** Ingredients and chemical composition of basal diets (As fed basis).

| Items | Diets |  |  |
| --- | --- | --- | --- |
|  | Starter | Grower | Finisher |
| Ingredients (gram/100 gram, as fed) |  |  |  |
| Yellow Corn | 53.4 | 58.0 | 61.3 |
| Soybean meal (44%) | 39.0 | 35.0 | 31.0 |
| Dicalcium phosphate | 1.7 | 1.5 | 1.3 |
| CaCO<sub>3</sub> (38%) | 1.7 | 1.4 | 1.3 |
| Sunflower oil | 3.0 | 3.0 | 4.0 |
| Sodium chloride | 0.3 | 0.3 | 0.3 |
| Methionine | 0.2 | 0.15 | 0.15 |
| Lysine | 0.2 | 0.15 | 0.15 |
| Premix* | 0.5 | 0.5 | 0.5 |
| Chemical composition |  |  |  |
| ME (kcal/kg) | 2900 | 3000 | 3100 |
| Crude protein (%) | 22 | 20.5 | 19 |
| Calcium (%) | 1.05 | 0.9 | 0.8 |
| Available Phosphorus (%) | 0.5 | 0.45 | 0.4 |
| Sodium (%) | 0.19 | 0.19 | 0.19 |
| Lysine (%) | 1.31 | 1.25 | 1.14 |
| Methionine (%) | 0.54 | 0.48 | 0.44 |
| Threonine (%) | 0.85 | 0.81 | 0.75 |
| Tryptophan (%) | 0.25 | 0.22 | 0.2 |

\* Vitamin and mineral content per kilogram of premix: vitamin A: 3,600,000 IU; vitamin D3: 800,000 IU; vitamin E: 7,200 IU; vitamin K3: 0.8 g; vitamin B1: 0.71 g; vitamin B2: 2.64 g; vitamin B3: 3.92 g, vitamin B5: 11.88 g; vitamin B6: 1.176 g; vitamin B12: 6 mg; folic acid: 0.4 g; biotin: 40 mg; choline chloride: 100 g; selenium: 80 mg; cobalt: 100 mg; iodine: 396 mg; copper: 4 g; zinc: 33.88 g; iron: 20 g; manganese: 39.68 g.

**Figure 1.**
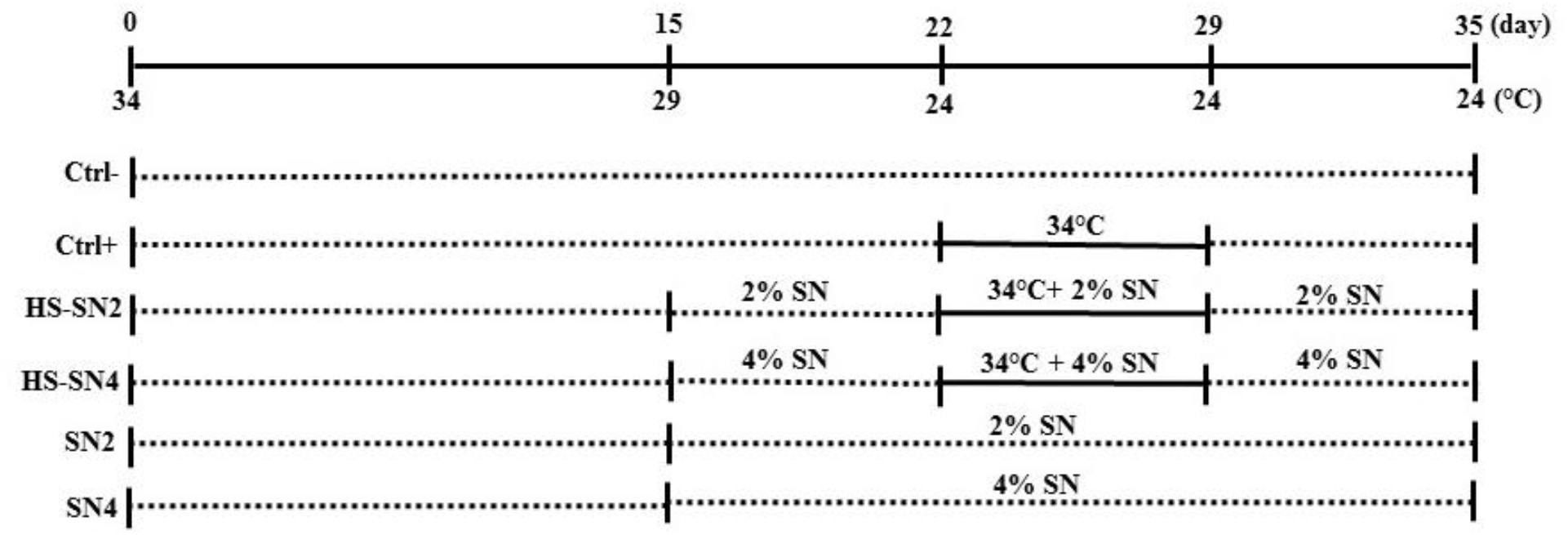
Experimental schedule of chronic heat exposure method. Ctrl-: negative control; Ctrl+: chronic heat-stressed broiler; HS-SN2: chronic heat-stressed broiler fed by 2% dietary stinging nettle; HS-SN4: chronic heat-stressed broilers fed by 4% stinging nettle; SN2: No exposure to chronic heat stress and fed by 2% dietary stinging nettle; SN4: No exposure to chronic heat stress and fed by 4% dietary stinging nettle; SN: stinting nettle.

### Dietary stinging nettle supplementation

All the birds were fed the basal diet from days 1 to 14. Subsequently, from days 14 to 35, stinging nettle powder was added to the basal diet in the treatment groups (HS-SN2, HS-SN4, SN2, and SN4). The stinging nettle was purchased from a local medicinal herb shop (Shiraz, Iran) and identified by a botanist (Shiraz University, School of Agriculture). The qualitative and quantitative analyses of several active compounds in stinging nettle powder was performed via gas chromatography–mass spectrometry (GC-MS) using Agilent 7890B GC system with Agilent 5977 Mass Selective Detector) (Table 2).

**Table 2.** Composition of the essential oils of used stinging nettle in the experiment (%).

| Essential oils | Content |
| --- | --- |
| Carvacrol | 37.89 |
| Naphthalene | 8.9 |
| Carvone | 8.39 |
| Butyldiene phthalide | 6.1 |
| Anethol | 3.8 |
| Phytol | 3.31 |
| Hexahydrofarnesyl acetone | 3.13 |
| Linalool | 2.51 |
| (E)-Geranyl acetone | 2.49 |
| Furanone | 2.45 |
| Caryophyllene oxide | 2.19 |
| Cumin aldehyde | 2.23 |
| β-caryophyllene | 1.87 |
| Bisabolene | 1.87 |
| Pentyl benzene | 1.65 |
| Cadinene | 1.64 |
| Limonene | 1.50 |
| Methyl chavicol | 1.42 |
| Kessane | 1.15 |
| Eugenol | 0.98 |
| Nonanal | 0.86 |
| Calamenene | 0.81 |
| β-selinene | 0.76 |
| α-terpineol | 0.56 |
| Cadina | 0.48 |
| Linalyl acetate | 0.46 |
| 2-pentyl furan | 0.33 |
| Copaene | 0.27 |

### Thermal programs and heat stress induction

On day zero, the ambient temperature was set as 34±1°C and did not change for the first 3 days. Afterwards, the room temperature gradually decreased 1°C every 2 days until reaching 24±1°C, and maintained at this temperature by controlling heat and ventilation until day 22. The cyclic heat stress was induced employing heater from days 22 to 29, and the relative humidity were kept at 50%±5% during this period (Altan et al., 2003). On day 22, the ambient temperature suddenly increased from 33±1°C to 38±1°C in the heat-stressed groups (HS, HS-SN2, and HS-SN4). The heat-stressed groups were reared in the controlled room at 38±1°C for 8 h/D (between 09:00 a.m. and 5:00 p.m.), followed by 24°C for the remaining 16 h/D. The NC, SN2, and SN4 groups were kept in separated rooms under normal temperature (24±1°C) for 24 h/D. During the heat stress period, the fans were kept active to maintain the homogeneity of the environmental temperature in room space. On day 29, the room temperature suddenly decreased from 38±1°C to 24±1°C in the heat-stressed groups, and from days 29 to 35, all the treatment groups were kept under ambient temperature (24±1°C).

### Growth performance

Their body weights (BW) were individually recorded on days 0, 14, 21, 29, and 35 to calculate the weight gain and average daily gain (ADG). Daily feed consumption was recorded per cage to determine the average daily feed intake (ADFI) and feed conversion ratio (FCR) in four periods (days 1-14, 14-29, 30-35, and 1-35). FI was calculated by subtracting the feed residues in the feeder from feed offered to each group in same periods. FCR was calculated per cage, and as the feed intake (g) divided by weight gain (g). At the end of the experiment (day 35), total weight gain, total feed intake, and total FCR were calculated in each group. Throughout the experiment, daily health status and mortality were recorded in each group.

### Clinical parameters

Rectal temperature was measured as an indicator of the core body temperature in broilers; in this regard, a digital thermometer (Pen-Type digital thermometer^®^, Dt-K101A, Shanghai, China) was inserted to a minimum depth of 3 cm in the cloaca, and the temperature of cloacal wall was recorded. During the heat stress exposure, the rectal temperature was recorded in two chickens per replicate (8 birds in per treatment group) starting about 4 h after the exposure to heat stress on days 22, 25, and 29. Panting frequency was also randomly measured based on video recordings on days 22, 25, and 29 (starting about 4 h after heat stress exposure) in two birds per replicate (8 birds in per treatment group). The panting frequency was calculated based on the number of breaths per minute.

### Blood and tissue samplings

On days 14, 21, 29, and 35, after 12-h fasting, three broilers from each replicate (12 birds in per treatment group) were randomly selected, weighed, and blood sampling (4 ml) was performed from the wing vein into the plain tubes. The blood samples were centrifuged at 750×g for 10 min at room temperature to separate sera. The collected sera were stored at -22°C for further analysis. On days 29 and 35, two birds from each replicate (8 samples from each treatment group) were randomly selected, slaughtered by cervical dislocation, and exsanguinated to get liver and intestine tissues. 2 cm-piece of right lob of liver and 3 cm of the middle portion of jejunum (from the entry of the ducts to Meckel’s diverticulum) were harvested and gently flushed with PBS to remove the gut contents. Subsequently, the intestinal and hepatic samples were placed into appropriate tubes, snap-frozen, and stored at -80°C until further analysis.

### Oxidative stress indices and HSP70 assays

The levels of oxidative stress biomarkers were analyzed in the collected sera, intestine, and liver samples. The serum contents of the total antioxidant capacity (T-AOC), malondialdehyde (MDA), reduced glutathione (GSH), activities of superoxide dismutase (SOD), and glutathione peroxidase (GSH-Px) were measured utilizing commercial kits (Kiazist Life Sciences, Hamedan, Iran) based on the provided instructions. The MDA and T-AOC levels were evaluated in liver and intestine samples with the commercial kits, according to the manufacturer’s instructions (Kiazist Life Sciences, Hamedan, Iran). The HSP70 levels of the collected hepatic and intestinal samples were determined using ELISA kit, according to the manufacturer’s instructions (Bioassay Technology Laboratory, Shanghai, China).

### Statistical analyses

Statistical analyses were carried out using the SPSS (SPSS for Windows, Version 22, SPSS Inc, Chicago, Illinois). All the data were subjected to one-way analysis of variance (ANOVA) followed by Post-Hoc Tukey HSD test. The results are expressed as the mean and standard error (mean±SE), and *P*<0.05 as a criterion of statistical significance.

## Results

Table 3 presents the effects of dietary stinging nettle on circulating oxidative stress indices of various treatments. There were no significant differences among some indices, on days 14 and 21, between all the studied groups (Table 3; *P*>0.05). Heat stress led to increased MDA levels and decreased T-AOC, SOD, and GSH-Px values in serum, liver, and intestine in the HS group compared to the control on days 29 and 35 (Table 3 and 4; *P*<0.05). Stinting nettle at both levels of 2% and 4% increased T-AOC, SOD, and GSH-Px, and decreased MDA levels in serum and various tissues in the heat-stressed broiler fed by supplemented diet (the HS-SN2 and HS-SN4 groups) and these effects was stranger in levels of 4% on days 29 and 35 (Table 3 and 4; *P*<0.05). In addition, Table 3 indicates that the serum activity of GSH was not significantly influenced by heat stress or stinting nettle at various treatments (*P*>0.05). Supplementing diet with stinting nettle in SN2 and SN4 groups increased the circulating T-AOC activity and decreased the circulating MDA on days 29 and 35.

**Table 3.**
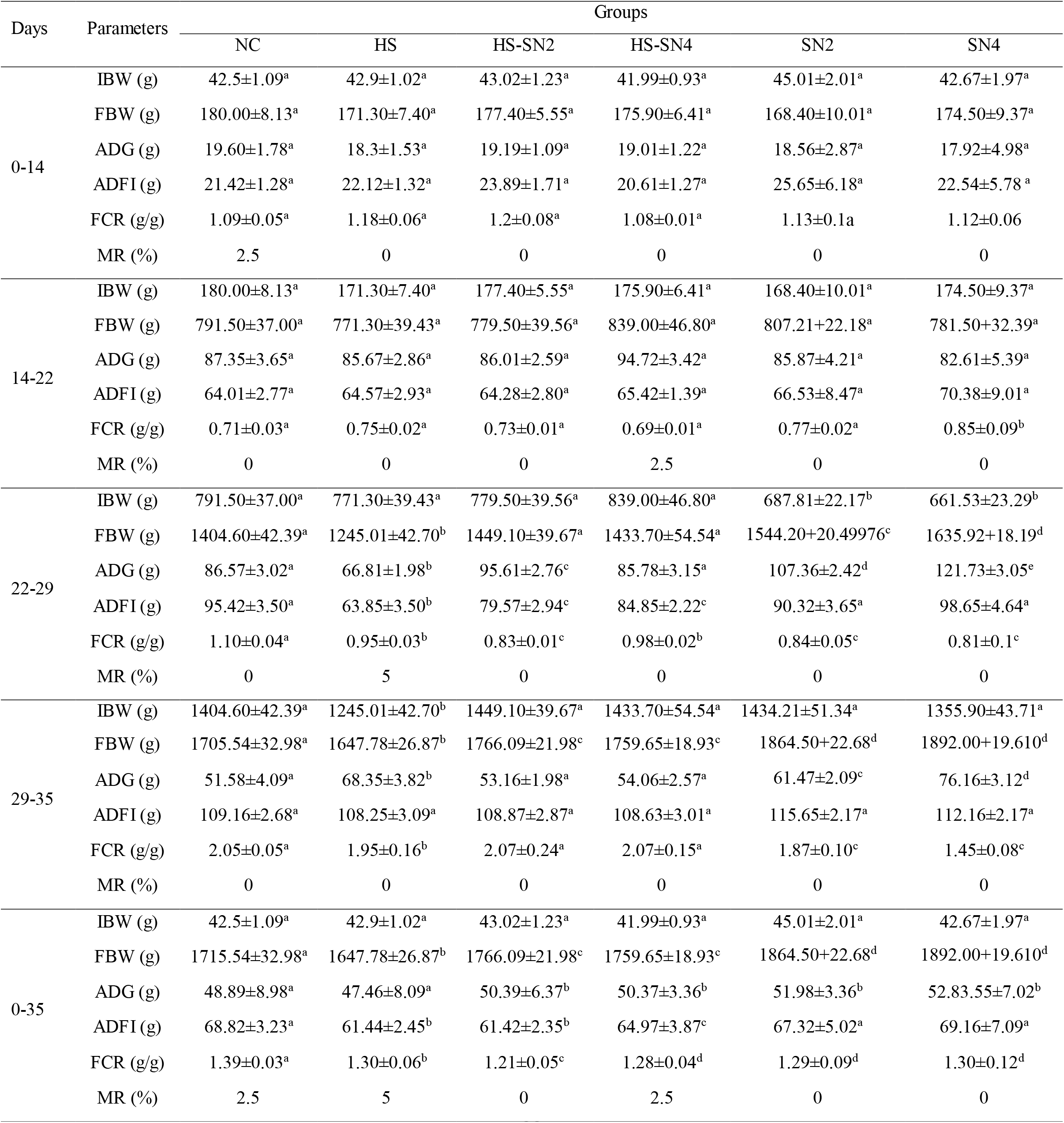
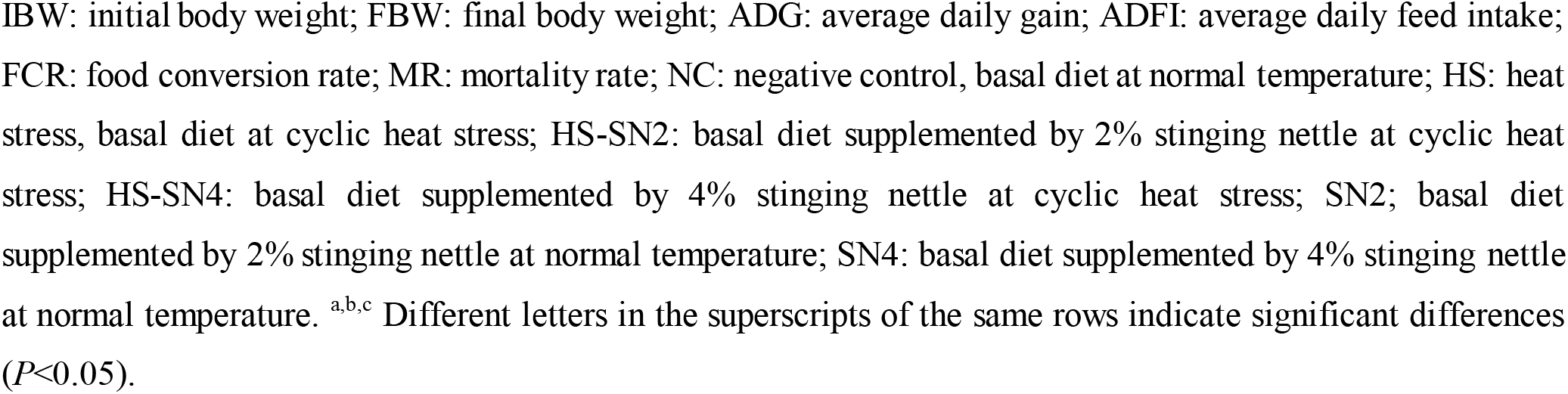
Effects of dietary stinging nettle on growth performance parameters (mean±SE) in various groups of broilers (n=40) subjected to heat stress.

| Days | Parameters | Groups |  |  |  |  |  |
| --- | --- | --- | --- | --- | --- | --- | --- |
|  |  | NC | HS | HS-SN2 | HS-SN4 | SN2 | SN4 |
| 0-14 | IBW (g) | 42.5±1.09<sup>a</sup> | 42.9±1.02<sup>a</sup> | 43.02±1.23<sup>a</sup> | 41.99±0.93<sup>a</sup> | 45.01±2.01<sup>a</sup> | 42.67±1.97<sup>a</sup> |
|  | FBW (g) | 180.00±8.13<sup>a</sup> | 171.30±7.40<sup>a</sup> | 177.40±5.55<sup>a</sup> | 175.90±6.41<sup>a</sup> | 168.40±10.01<sup>a</sup> | 174.50±9.37<sup>a</sup> |
|  | ADG (g) | 19.60±1.78<sup>a</sup> | 18.3±1.53<sup>a</sup> | 19.19±1.09<sup>a</sup> | 19.01±1.22<sup>a</sup> | 18.56±2.87<sup>a</sup> | 17.92±4.98<sup>a</sup> |
|  | ADFI (g) | 21.42±1.28<sup>a</sup> | 22.12±1.32<sup>a</sup> | 23.89±1.71<sup>a</sup> | 20.61±1.27<sup>a</sup> | 25.65±6.18<sup>a</sup> | 22.54±5.78<sup>a</sup> |
|  | FCR (g/g) | 1.09±0.05<sup>a</sup> | 1.18±0.06<sup>a</sup> | 1.2±0.08<sup>a</sup> | 1.08±0.01<sup>a</sup> | 1.13±0.1a | 1.12±0.06 |
|  | MR (%) | 2.5 | 0 | 0 | 0 | 0 | 0 |
| 14-22 | IBW (g) | 180.00±8.13<sup>a</sup> | 171.30±7.40<sup>a</sup> | 177.40±5.55<sup>a</sup> | 175.90±6.41<sup>a</sup> | 168.40±10.01<sup>a</sup> | 174.50±9.37<sup>a</sup> |
|  | FBW (g) | 791.50±37.00<sup>a</sup> | 771.30±39.43<sup>a</sup> | 779.50±39.56<sup>a</sup> | 839.00±46.80<sup>a</sup> | 807.21±22.18<sup>a</sup> | 781.50±32.39<sup>a</sup> |
|  | ADG (g) | 87.35±3.65<sup>a</sup> | 85.67±2.86<sup>a</sup> | 86.01±2.59<sup>a</sup> | 94.72±3.42<sup>a</sup> | 85.87±4.21<sup>a</sup> | 82.61±5.39<sup>a</sup> |
|  | ADFI (g) | 64.01±2.77<sup>a</sup> | 64.57±2.93<sup>a</sup> | 64.28±2.80<sup>a</sup> | 65.42±1.39<sup>a</sup> | 66.53±8.47<sup>a</sup> | 70.38±9.01<sup>a</sup> |
|  | FCR (g/g) | 0.71±0.03<sup>a</sup> | 0.75±0.02<sup>a</sup> | 0.73±0.01<sup>a</sup> | 0.69±0.01<sup>a</sup> | 0.77±0.02<sup>a</sup> | 0.85±0.09<sup>b</sup> |
|  | MR (%) | 0 | 0 | 0 | 2.5 | 0 | 0 |
| 22-29 | IBW (g) | 791.50±37.00<sup>a</sup> | 771.30±39.43<sup>a</sup> | 779.50±39.56<sup>a</sup> | 839.00±46.80<sup>a</sup> | 687.81±22.17<sup>b</sup> | 661.53±23.29<sup>b</sup> |
|  | FBW (g) | 1404.60±42.39<sup>a</sup> | 1245.01±42.70<sup>b</sup> | 1449.10±39.67<sup>a</sup> | 1433.70±54.54<sup>a</sup> | 1544.20±20.49976<sup>c</sup> | 1635.92±18.19<sup>d</sup> |
|  | ADG (g) | 86.57±3.02<sup>a</sup> | 66.81±1.98<sup>b</sup> | 95.61±2.76<sup>c</sup> | 85.78±3.15<sup>a</sup> | 107.36±2.42<sup>d</sup> | 121.73±3.05<sup>e</sup> |
|  | ADFI (g) | 95.42±3.50<sup>a</sup> | 63.85±3.50<sup>b</sup> | 79.57±2.94<sup>c</sup> | 84.85±2.22<sup>c</sup> | 90.32±3.65<sup>a</sup> | 98.65±4.64<sup>a</sup> |
|  | FCR (g/g) | 1.10±0.04<sup>a</sup> | 0.95±0.03<sup>b</sup> | 0.83±0.01<sup>c</sup> | 0.98±0.02<sup>b</sup> | 0.84±0.05<sup>c</sup> | 0.81±0.1<sup>c</sup> |
|  | MR (%) | 0 | 5 | 0 | 0 | 0 | 0 |
| 29-35 | IBW (g) | 1404.60±42.39<sup>a</sup> | 1245.01±42.70<sup>b</sup> | 1449.10±39.67<sup>a</sup> | 1433.70±54.54<sup>a</sup> | 1434.21±51.34<sup>a</sup> | 1355.90±43.71<sup>a</sup> |
|  | FBW (g) | 1705.54±32.98<sup>a</sup> | 1647.78±26.87<sup>b</sup> | 1766.09±21.98<sup>c</sup> | 1759.65±18.93<sup>c</sup> | 1864.50±22.68<sup>d</sup> | 1892.00±19.610<sup>d</sup> |
|  | ADG (g) | 51.58±4.09<sup>a</sup> | 68.35±3.82<sup>b</sup> | 53.16±1.98<sup>a</sup> | 54.06±2.57<sup>a</sup> | 61.47±2.09<sup>c</sup> | 76.16±3.12<sup>d</sup> |
|  | ADFI (g) | 109.16±2.68<sup>a</sup> | 108.25±3.09<sup>a</sup> | 108.87±2.87<sup>a</sup> | 108.63±3.01<sup>a</sup> | 115.65±2.17<sup>a</sup> | 112.16±2.17<sup>a</sup> |
|  | FCR (g/g) | 2.05±0.05<sup>a</sup> | 1.95±0.16<sup>b</sup> | 2.07±0.24<sup>a</sup> | 2.07±0.15<sup>a</sup> | 1.87±0.10<sup>c</sup> | 1.45±0.08<sup>c</sup> |
|  | MR (%) | 0 | 0 | 0 | 0 | 0 | 0 |
| 0-35 | IBW (g) | 42.5±1.09<sup>a</sup> | 42.9±1.02<sup>a</sup> | 43.02±1.23<sup>a</sup> | 41.99±0.93<sup>a</sup> | 45.01±2.01<sup>a</sup> | 42.67±1.97<sup>a</sup> |
|  | FBW (g) | 1715.54±32.98<sup>a</sup> | 1647.78±26.87<sup>b</sup> | 1766.09±21.98<sup>c</sup> | 1759.65±18.93<sup>c</sup> | 1864.50±22.68<sup>d</sup> | 1892.00±19.610<sup>d</sup> |
|  | ADG (g) | 48.89±8.98<sup>a</sup> | 47.46±8.09<sup>a</sup> | 50.39±6.37<sup>b</sup> | 50.37±3.36<sup>b</sup> | 51.98±3.36<sup>b</sup> | 52.83.55±7.02<sup>b</sup> |
|  | ADFI (g) | 68.82±3.23<sup>a</sup> | 61.44±2.45<sup>b</sup> | 61.42±2.35<sup>b</sup> | 64.97±3.87<sup>c</sup> | 67.32±5.02<sup>a</sup> | 69.16±7.09<sup>a</sup> |
|  | FCR (g/g) | 1.39±0.03<sup>a</sup> | 1.30±0.06<sup>b</sup> | 1.21±0.05<sup>c</sup> | 1.28±0.04<sup>d</sup> | 1.29±0.09<sup>d</sup> | 1.30±0.12<sup>d</sup> |
|  | MR (%) | 2.5 | 5 | 0 | 2.5 | 0 | 0 |

IBW: initial body weight; FBW: final body weight; ADG: average daily gain; ADFI: average daily feed intake; FCR: food conversion rate; MR: mortality rate; NC: negative control, basal diet at normal temperature; HS: heat stress, basal diet at cyclic heat stress; HS-SN2: basal diet supplemented by 2% stinging nettle at cyclic heat stress; HS-SN4: basal diet supplemented by 4% stinging nettle at cyclic heat stress; SN2: basal diet supplemented by 2% stinging nettle at normal temperature; SN4: basal diet supplemented by 4% stinging nettle at normal temperature. <sup>a,b,c</sup> Different letters in the superscripts of the same rows indicate significant differences ( $P<0.05$ ).

**Table 4.** Effect of two different levels of dietary stinging nettle on rectal temperature and panting frequency (mean±SE) of broilers subjected to heat stress.

| Parameters | Days | Groups |  |  |  |  |  |
| --- | --- | --- | --- | --- | --- | --- | --- |
|  |  | NC | HS | HS-SN2 | HS-SN4 | SN2 | SN4 |
| Panting frequency (#/min) | 22 | 80.10±2.12<sup>a</sup> | 142.90±2.46<sup>b</sup> | 140.10±0.92<sup>b,c</sup> | 137.10±1.16<sup>b</sup> | 85.54±1.36<sup>a</sup> | 79.05±1.66<sup>a</sup> |
|  | 25 | 80.20±2.09<sup>a</sup> | 141.90±1.17<sup>b</sup> | 138.70±1.73<sup>b</sup> | 139.50±1.65<sup>b</sup> | 79.98± 3.09<sup>a</sup> | 87.11±3.07<sup>a</sup> |
|  | 29 | 79.90±2.05<sup>a</sup> | 136.50±1.06<sup>b</sup> | 136.80±1.29<sup>b</sup> | 138.01±1.58<sup>b</sup> | 86.42±2.09<sup>a</sup> | 90.76±2.10<sup>a</sup> |
| Rectal temperature (°C) | 22 | 40.47±0.15<sup>a</sup> | 43.25±0.17<sup>b</sup> | 43.30±0.15<sup>b</sup> | 42.89±0.18<sup>b</sup> | 41.01±0.09<sup>a</sup> | 41.32±0.21<sup>a</sup> |
|  | 25 | 40.78±0.52<sup>a</sup> | 43.25±0.17<sup>b</sup> | 43.89±0.17<sup>b</sup> | 42.89±0.18<sup>b</sup> | 40.09±0.04<sup>a</sup> | 40.41±0.35<sup>a</sup> |
|  | 29 | 40.75±0.20<sup>a</sup> | 42.87±0.17<sup>b</sup> | 42.53±0.15<sup>b</sup> | 42.80±0.20<sup>b</sup> | 40.12±0.12<sup>a</sup> | 40.28±0.29<sup>a</sup> |

NC: negative control, basal diet at normal temperature; HS: heat stress, basal diet at cyclic heat stress; HS-SN2: basal diet supplemented by 2% stinging nettle at cyclic heat stress; HS-SN4: basal diet supplemented by 4% stinging nettle at cyclic heat stress; SN2: basal diet supplemented by 2% stinging nettle at normal temperature; SN4: basal diet supplemented by 4% stinging nettle at normal temperature.
<sup>a,b,c</sup> Different letters in the superscripts of the same rows indicate significant differences ( $P<0.05$ ).

As could be seen in Table 4, HSP70 levels in the liver and intestine increased in the heat-stressed groups compared to the NC on days 29 and 35 (*P*<0.05). The hepatic and intestinal levels of HSP70 were significantly lower in the heat-stressed broilers fed with 2% and 4% stinging nettle compared to the heat-stressed birds on the basal diet (Table 4; *P*<0.05).

Table 5 displays an overview of the growth performance in the heat-stressed broilers fed with different levels of stinging nettle. Heat stress caused decreased BW, ADG, and ADFI in the HS compared to the NC (*P*<0.05). Receiving stinting nettle at 2% and 4% levels in the broilers exposed to heat stress triggered a significant increase in the BW, ADG, and ADFI compared to those fed by basal diet on day 29 and 35 (Table 5; *P*<0.05). The lowest final weigh gain and highest mortality rate belonged to the heat-stressed group compared to the other treatments on days 35 (P<0.05).

**Table 5.**
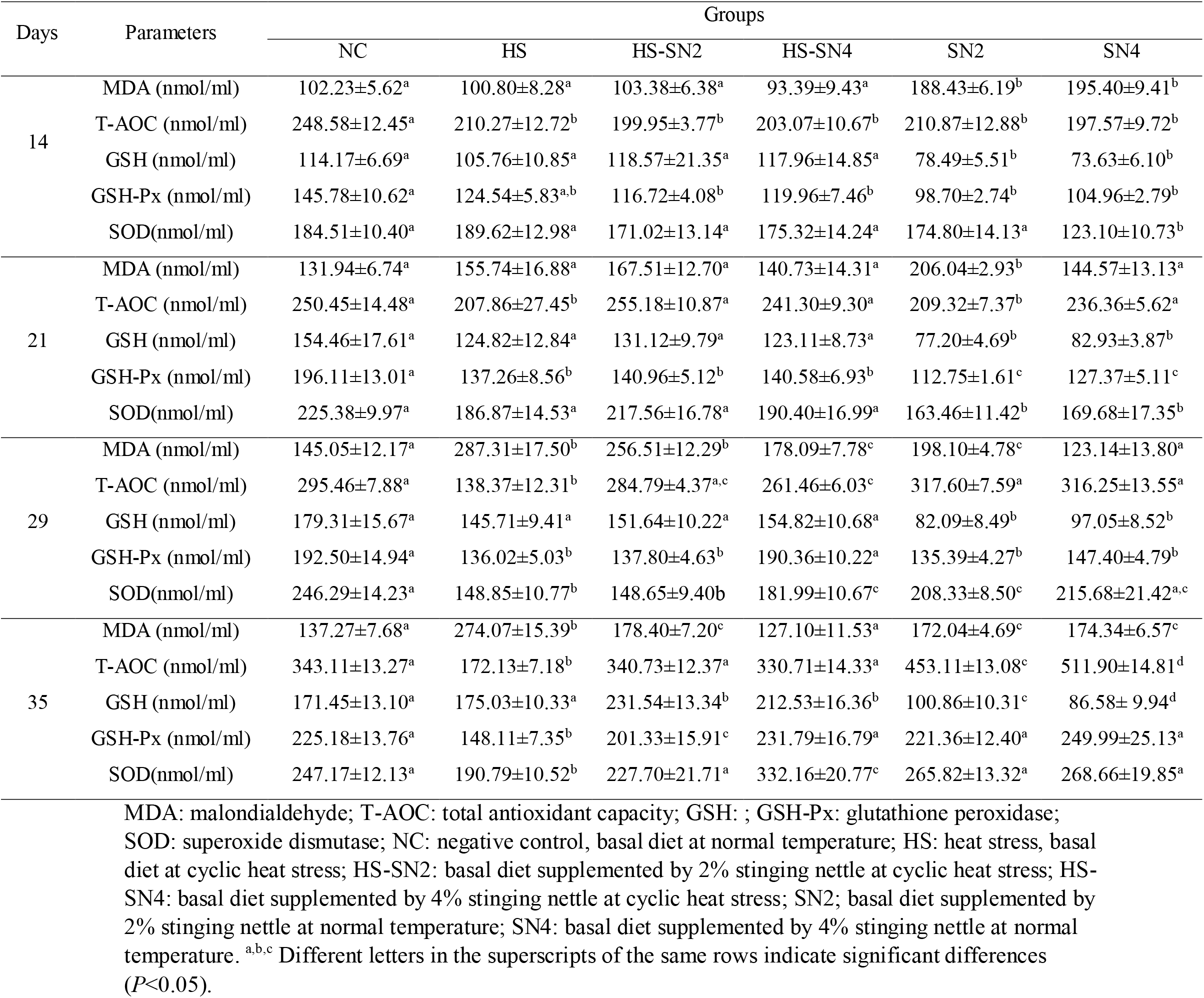
Effects of dietary stinging nettle on serum oxidative stress indices (mean±SE) in various groups of broilers (n=40) subjected to heat stress.

| Days | Parameters | Groups |  |  |  |  |  |
| --- | --- | --- | --- | --- | --- | --- | --- |
|  |  | NC | HS | HS-SN2 | HS-SN4 | SN2 | SN4 |
| 14 | MDA (nmol/ml) | 102.23±5.62<sup>a</sup> | 100.80±8.28<sup>a</sup> | 103.38±6.38<sup>a</sup> | 93.39±9.43<sup>a</sup> | 188.43±6.19<sup>b</sup> | 195.40±9.41<sup>b</sup> |
|  | T-AOC (nmol/ml) | 248.58±12.45<sup>a</sup> | 210.27±12.72<sup>b</sup> | 199.95±3.77<sup>b</sup> | 203.07±10.67<sup>b</sup> | 210.87±12.88<sup>b</sup> | 197.57±9.72<sup>b</sup> |
|  | GSH (nmol/ml) | 114.17±6.69<sup>a</sup> | 105.76±10.85<sup>a</sup> | 118.57±21.35<sup>a</sup> | 117.96±14.85<sup>a</sup> | 78.49±5.51<sup>b</sup> | 73.63±6.10<sup>b</sup> |
|  | GSH-Px (nmol/ml) | 145.78±10.62<sup>a</sup> | 124.54±5.83<sup>a,b</sup> | 116.72±4.08<sup>b</sup> | 119.96±7.46<sup>b</sup> | 98.70±2.74<sup>b</sup> | 104.96±2.79<sup>b</sup> |
|  | SOD(nmol/ml) | 184.51±10.40<sup>a</sup> | 189.62±12.98<sup>a</sup> | 171.02±13.14<sup>a</sup> | 175.32±14.24<sup>a</sup> | 174.80±14.13<sup>a</sup> | 123.10±10.73<sup>b</sup> |
| 21 | MDA (nmol/ml) | 131.94±6.74<sup>a</sup> | 155.74±16.88<sup>a</sup> | 167.51±12.70<sup>a</sup> | 140.73±14.31<sup>a</sup> | 206.04±2.93<sup>b</sup> | 144.57±13.13<sup>a</sup> |
|  | T-AOC (nmol/ml) | 250.45±14.48<sup>a</sup> | 207.86±27.45<sup>b</sup> | 255.18±10.87<sup>a</sup> | 241.30±9.30<sup>a</sup> | 209.32±7.37<sup>b</sup> | 236.36±5.62<sup>a</sup> |
|  | GSH (nmol/ml) | 154.46±17.61<sup>a</sup> | 124.82±12.84<sup>a</sup> | 131.12±9.79<sup>a</sup> | 123.11±8.73<sup>a</sup> | 77.20±4.69<sup>b</sup> | 82.93±3.87<sup>b</sup> |
|  | GSH-Px (nmol/ml) | 196.11±13.01<sup>a</sup> | 137.26±8.56<sup>b</sup> | 140.96±5.12<sup>b</sup> | 140.58±6.93<sup>b</sup> | 112.75±1.61<sup>c</sup> | 127.37±5.11<sup>c</sup> |
|  | SOD(nmol/ml) | 225.38±9.97<sup>a</sup> | 186.87±14.53<sup>a</sup> | 217.56±16.78<sup>a</sup> | 190.40±16.99<sup>a</sup> | 163.46±11.42<sup>b</sup> | 169.68±17.35<sup>b</sup> |
| 29 | MDA (nmol/ml) | 145.05±12.17<sup>a</sup> | 287.31±17.50<sup>b</sup> | 256.51±12.29<sup>b</sup> | 178.09±7.78<sup>c</sup> | 198.10±4.78<sup>c</sup> | 123.14±13.80<sup>a</sup> |
|  | T-AOC (nmol/ml) | 295.46±7.88<sup>a</sup> | 138.37±12.31<sup>b</sup> | 284.79±4.37<sup>a,c</sup> | 261.46±6.03<sup>c</sup> | 317.60±7.59<sup>a</sup> | 316.25±13.55<sup>a</sup> |
|  | GSH (nmol/ml) | 179.31±15.67<sup>a</sup> | 145.71±9.41<sup>a</sup> | 151.64±10.22<sup>a</sup> | 154.82±10.68<sup>a</sup> | 82.09±8.49<sup>b</sup> | 97.05±8.52<sup>b</sup> |
|  | GSH-Px (nmol/ml) | 192.50±14.94<sup>a</sup> | 136.02±5.03<sup>b</sup> | 137.80±4.63<sup>b</sup> | 190.36±10.22<sup>a</sup> | 135.39±4.27<sup>b</sup> | 147.40±4.79<sup>b</sup> |
|  | SOD(nmol/ml) | 246.29±14.23<sup>a</sup> | 148.85±10.77<sup>b</sup> | 148.65±9.40<sup>b</sup> | 181.99±10.67<sup>c</sup> | 208.33±8.50<sup>c</sup> | 215.68±21.42<sup>a,c</sup> |
| 35 | MDA (nmol/ml) | 137.27±7.68<sup>a</sup> | 274.07±15.39<sup>b</sup> | 178.40±7.20<sup>c</sup> | 127.10±11.53<sup>a</sup> | 172.04±4.69<sup>c</sup> | 174.34±6.57<sup>c</sup> |
|  | T-AOC (nmol/ml) | 343.11±13.27<sup>a</sup> | 172.13±7.18<sup>b</sup> | 340.73±12.37<sup>a</sup> | 330.71±14.33<sup>a</sup> | 453.11±13.08<sup>c</sup> | 511.90±14.81<sup>d</sup> |
|  | GSH (nmol/ml) | 171.45±13.10<sup>a</sup> | 175.03±10.33<sup>a</sup> | 231.54±13.34<sup>b</sup> | 212.53±16.36<sup>b</sup> | 100.86±10.31<sup>c</sup> | 86.58± 9.94<sup>d</sup> |
|  | GSH-Px (nmol/ml) | 225.18±13.76<sup>a</sup> | 148.11±7.35<sup>b</sup> | 201.33±15.91<sup>c</sup> | 231.79±16.79<sup>a</sup> | 221.36±12.40<sup>a</sup> | 249.99±25.13<sup>a</sup> |
|  | SOD(nmol/ml) | 247.17±12.13<sup>a</sup> | 190.79±10.52<sup>b</sup> | 227.70±21.71<sup>a</sup> | 332.16±20.77<sup>c</sup> | 265.82±13.32<sup>a</sup> | 268.66±19.85<sup>a</sup> |

MDA: malondialdehyde; T-AOC: total antioxidant capacity; GSH: ; GSH-Px: glutathione peroxidase; SOD: superoxide dismutase; NC: negative control, basal diet at normal temperature; HS: heat stress, basal diet at cyclic heat stress; HS-SN2: basal diet supplemented by 2% stinging nettle at cyclic heat stress; HS-SN4: basal diet supplemented by 4% stinging nettle at cyclic heat stress; SN2; basal diet supplemented by 2% stinging nettle at normal temperature; SN4: basal diet supplemented by 4% stinging nettle at normal temperature. <sup>a,b,c</sup> Different letters in the superscripts of the same rows indicate significant differences ( $P<0.05$ ).

Table 6 shows the effects of stinging nettle supplementation on rectal temperature and panting of broilers subjected to heat stress. All the groups exposed to heat stress had a significantly higher rectal temperature and panting frequency compared to the treatments on normal temperature on days 22, 25, and 29 (P<0.05). Feeding with stinting nettle in the HS-SN2 and HS-SN4 had no significant effects on rectal temperature and panting frequency at different times of experiment (P>0.05).

**Table 6.** Effects of dietary stinging nettle on liver and intestine levels of oxidative stress indices and HSP70 (mean±SE) in various groups of broilers (n=40) subjected to heat stress.

| Days | Tissues | Parameters | Groups |  |  |  |  |  |
| --- | --- | --- | --- | --- | --- | --- | --- | --- |
|  |  |  | NC | HS | HS-SN2 | HS-SN4 | SN2 | SN4 |
| 29 | Liver | MDA (nmol/ml) | 120.50±8.91<sup>a</sup> | 381.21±27.14<sup>b</sup> | 312.72±25.42<sup>c</sup> | 179.08±20.12<sup>a</sup> | 128.43±11.65<sup>a</sup> | 129.77±15.14<sup>a</sup> |
|  |  | T-AOC(nmol/ml) | 394.23±23.36<sup>a</sup> | 309.11±20.72<sup>b</sup> | 330.60±12.23<sup>b</sup> | 349.02±16.62<sup>b</sup> | 512.06±21.60<sup>c</sup> | 482.81±26.24<sup>c</sup> |
|  |  | HSP70 (ng/ml) | 24.08±1.49<sup>a,c</sup> | 58.08±5.46<sup>b</sup> | 33.14±3.71<sup>a</sup> | 14.25±1.92<sup>c</sup> | 26.08±2.46<sup>a</sup> | 21.11±2.20<sup>a,c</sup> |
|  | Intestine | MDA (nmol/ml) | 124.49±29.81<sup>a</sup> | 444.26±17.74<sup>b</sup> | 348.19±25.81<sup>c</sup> | 231.55±22.86<sup>d</sup> | 133.23±10.78<sup>a</sup> | 126.30±12.87<sup>a</sup> |
|  |  | T-AOC(nmol/ml) | 321.42±13.12<sup>a</sup> | 234.08±16.32<sup>b</sup> | 270.06±12.20<sup>c</sup> | 281.87±10.27<sup>c</sup> | 492.87±15.15<sup>d</sup> | 546.44±14.13<sup>e</sup> |
|  |  | HSP70 (ng/ml) | 18.65±1.73<sup>a</sup> | 67.72±1.91<sup>b</sup> | 45.71±3.91<sup>c</sup> | 25.19±2.49<sup>a</sup> | 23.91±2.64<sup>a</sup> | 24.19±3.64<sup>a</sup> |
| 35 | Liver | MDA (nmol/ml) | 126.04±8.89<sup>a</sup> | 324.58±18.56<sup>b</sup> | 202.09±20.20<sup>c</sup> | 134.42±26.23<sup>a</sup> | 140.31±10.83<sup>a</sup> | 137.58±9.08<sup>a</sup> |
|  |  | T-AOC(nmol/ml) | 404.15±22.29<sup>a</sup> | 340.18±17.95<sup>b</sup> | 407.78±15.16<sup>a</sup> | 390.58±16.18<sup>a</sup> | 527.72±14.50<sup>c</sup> | 583.07±16.89<sup>c</sup> |
|  |  | HSP70 (ng/ml) | 24.08±1.49<sup>a</sup> | 50.86±3.64<sup>b</sup> | 22.48±1.96<sup>a</sup> | 10.91±1.54<sup>c</sup> | 26.08±2.46<sup>a</sup> | 21.11±2.20<sup>a</sup> |
|  | Intestine | MDA (nmol/ml) | 95.28±12.14<sup>a</sup> | 363.81±20.33<sup>b</sup> | 276.56±15.93<sup>c</sup> | 192.66±14.79<sup>d</sup> | 113.26±13.08<sup>a</sup> | 100.44±11.05<sup>a</sup> |
|  |  | T-AOC(nmol/ml) | 354.22±18.43<sup>a</sup> | 323.51±13.79<sup>a</sup> | 337.26±19.39<sup>a</sup> | 332.06±33.26<sup>a</sup> | 613.29±16.66<sup>b</sup> | 560.96±17.67<sup>b</sup> |
|  |  | HSP70 (ng/ml) | 18.65±1.73<sup>a</sup> | 39.16±4.15<sup>b</sup> | 33.22±3.92<sup>a</sup> | 17.46±1.10<sup>a</sup> | 21.08±2.40<sup>a</sup> | 19.32±2.70<sup>a</sup> |

MDA: malondialdehyde; T-AOC: total antioxidant capacity; HSP70; heat shock protein 70; NC: negative control, basal diet at normal temperature; HS: heat stress, basal diet at cyclic heat stress; HS-SN2: basal diet supplemented by 2% stinging nettle at cyclic heat stress; HS-SN4: basal diet supplemented by 4% stinging nettle at cyclic heat stress; SN2; basal diet supplemented by 2% stinging nettle at normal temperature; SN4: basal diet supplemented by 4% stinging nettle at normal temperature. <sup>a,b,c</sup> Different letters in the superscripts of the same rows indicate significant differences ( $P<0.05$ ).

## Discussion

Heat stress leads to oxidative stress and multiple organ damages in broilers (Chen et al., 2020). Therefore, several researchers have emphasized employing managerial solutions to reduce harmful effects of heat stress conditions on these birds. Using feed additives and dietary supplementations with antioxidant materials and plants are common methods to obtain the best results. Hence, the present study evaluated the effects of dietary stinging nettle on heat stress-induced oxidative stress and organ damages in broilers. This study revealed that dietary supplementation of stinging nettle powder at 4% reduced the circulating and tissue oxidative stress indices of broilers subjected to heat stress (Table 3 and 4). Furthermore, this plant improved the growth performance in the heat-stressed broilers (Table 5). Accordingly, using stinging nettle could be suggested to combat the side effects of chronic heat stress in broiler production industry.

Stinging nettle has various compounds with antioxidant effects, including terpenoid phenol, flavonoids, alpha tochopherol, and ascorbic acid (Surai, 2014). The main terpenoids found in stinting nettle are carvacrol and carvone, which account for 46.28% of the oil in this plant (Table 2). Upton (2013) stated that carvacrol and carvone are of antioxidative, growth-promoting, antibacterial, and antiviral activities. Terpenoids and phenolic compounds in stinging nettle inhibit oxidative stress through various mechanisms, such as scavenging ROS, inhibiting lipid peroxidation, activating antioxidant enzymes, metal chelating activity, and increasing uric acid levels (Behrooj et al., 2012). The supplemented diets with stinging nettle caused a decrease in MDA and an increase in antioxidant content of serum, meat, lung, liver, and intestine in the broilers (Botsoglou et al., 2002; Loetcher et al. 2013a,b). In this regard, the effects of stinting nettle on certain oxidative stress indices, including MDA, T-AOC, SOD, GSH, and GSH-Px, which were assessed to evaluate the oxidative status in broilers subjected to heat stress (Table 3 and 4). Following heat exposure, the contents of MDA in serum (49%), liver (68%), and intestine (72%) in the HS group significantly increased compared to the NC on day 29; these results are consistent with previous studies stating that heat stress leads to overproduction of MDA (Abo Ghanima et al., 2020). In line with these results, Mujahid et al. (2007) stated that heat exposure resulted in peroxidation of lipids and higher mitochondrial MDA levels in cells. In the studied birds, both dietary levels of stinging nettle in the HS-SN2 and HS-SN4 groups resulted into the reduction of the serum, intestinal, and hepatic levels of MDA compared to the HS group fed by the basal diet on days 29 and 35; this effect was stranger at 4% stinging nettle supplementation (Table 3 and 4; *P*<0.05). In agreement with these results, Golchin et al. (2004) reported that water extract of stinting nettle (50 mg/ml) has a stronger effect on inhibiting lipid peroxidation compared to alpha tocopherol (60 mg/mL). They indicated that stinging nettle reduced lipid peroxidation by 39%, while alpha tocopherol showed 30% of inhibition. In addition, Ahmadipour and Khajali (2019) reported that stinging nettle powder at the level of 1.5% had a potent antioxidant activity on reducing lipid peroxidation and circulating MDA in the broilers exposed to pulmonary hypertension syndrome. Based on the current investigation, 4% dietary stinging nettle led to a significant decrease in the serum and tissue levels of MDA, indicating that this herb could attenuate the oxidative stress induced by heat exposure in broilers. Following the exposure to stressful conditions, the antioxidant enzymes (SOD and GSH-Px) act as the first line of body’s antioxidant defense to scavenge and neutralize the free radicals and maintain health status of cells. The exposure to chronic heat causes a decrease in antioxidant enzymes expression and activity. This reduction could be attributed to overconsumpation or thermal/oxidative inactivation of the enzymes (Moriyama-Gonda et al., 2002). As shown in Table 3, chronic heat stress in the HS group decreased significantly the serum levels of T-AOC, SOD, and GSH-Px activity compared to the NC on day 29. In accordance with these results, Nawab et al. (2019) stated that the levels of T-OAC, SOD, and GSH-Px reduced in heat-stressed broilers compared to those on ambient temperature. Dietary stinging nettle at 2% and 4% significantly increased T-AOC levels in serum, liver, and intestine of the HS-SN2, HS-SN4, SN2, and SN4 compared to the HS on days 29 and 35 (Table 3 and 4). The serum activity of GSH-Px and SOD increased in the HS-SN4 compared to the HS and HS-SN2 groups; there were no significant differences between the HS-SN4 and the NC on day 29. In line with our findings, Ahmadipour and Khajali (2019) reported that the stinging nettle powder increased the expression of antioxidant enzymes in lung and liver of diseased broilers. They indicated that stinging nettle reduced oxidative stress by reducing MDA production and increasing the activities of antioxidant enzymes in the broilers with pulmonary hypertension syndrome. As represented in Table 3 and 4, the oxidative status improved in the heat-stressed groups (HS, HS-SN2, and HS-SN4) on day 7 following the end of heat exposure, yet this effect was higher in the group fed by supplemented diet with 4% stinging nettle. These findings revealed that stinging nettle supplementation at level 4% had positive and significant effects on improving the antioxidant status of broilers during heat exposure and 7 days following the exposure to heat stress. It could be concluded that supplementing diet with 4% stinging nettle increased the activity levels of antioxidant enzymes that protect birds against adverse effects of oxidative stress following exposure to heat stress.

In response to stressful conditions, cellular proteins are oxidized and denatured. Following heat exposure, HSPs are synthesized to facilitate the folding and refolding of stress-denatured proteins and prevent proteins degradation and cells against lethal thermal damages (Ahmed et al. 2012). Previous studies have established that the hepatic and intestinal levels of HSP70 enhance the antioxidant capacity and inhibit the lipid peroxidation of chickens, and HSP70 protects the intestinal mucosa and hepatic cells against cellular injuries caused by heat stress (Surai et al., 2019). In the studied broilers, the hepatic and ileal concentrations of HSP70 significantly increased in the HS compared to the NC group on day 29 (Table 4, *P*<0.05). Gu et al. (2012) showed that HSP70 is synthesized in broilers under heat exposure condition and exerting protective effects against the toxic effects of heat stress. In the heat-stressed broilers, the expression of HSP70 increased the tissue dependently, and various tissues respond differently to the heat challenge (Rajkumar et al., 2017). In this study, ileal cells had significantly higher HSP70 levels compared to hepatic cells (Table 4). Under chronic heat stress, stinging nettle supplementation at both 2% and 4% levels caused a decrease in hepatic and intestinal contents of HSP70; this effect was more severe at 4%. There were no significant differences in HSP70 levels in the tissue samples between the HS-SN4 and NC groups on day 29 (Table 4; *P*>0.05). Furthermore, 7 days after the end of heat exposure (day 35), the hepatic and intestinal levels of HSP70 in the HS-SN2 and HS-SN4 were significantly lower compared to those of the HS group. In line with these results, Surai et al. (2019) stated that medicinal plants with a high antioxidant content reduced the HSP expression in the birds subjected to stressful conditions. However, there are limited reviews on the effects of stinging nettle on HSP70 levels in broilers under stressful conditions. The observations by the present study could support the hypothesis that dietary stinging nettle significantly reduces HSP70 synthesis in heat-stressed broilers. Therefore, the current experiment revealed that dietary stinging nettle had dose-dependent effects on oxidative stress. The present study shed light on the fact that dietary stinging nettle at level 4% is effective on the improvement of oxidative status in broilers and control of oxidative stress.

Growth performance, weight gain, and feed efficiency are the main economic parameters in poultry production and control of all the factors affecting these parameters is the most important goal during the rearing of commercial poultry (Rosa et al., 2007). Heat stress is one of the most important environmental disruptors with negative effects on food intake and weight gain in broilers (Quinteiro-Filho et al., 2010). Liu et al. (2014) showed that heat stress led to a decrease in the feed intake and weight gain in broilers. As seen in Table 5, on days 29 and 35, the heat-stressed broilers had significantly lower BW, ADG, ADFI, and higher FCR compared to those under normal temperature. Stinging nettle supplementation at both levels of 2% (HS-SN2 group) and 4% (HS-SN4 group) increased BW, ADG, and ADFI compared to the HS group on day 29; this effect was more potent in 2% stinging nettle supplementation (Table 5; *P*<0.05). Dietary stinging nettle had significant positive effects on BW, ADG, and FCR in the SN2 and SN4 groups compared to those fed by the basal diet on days 29 and 35 (*P*>0.05). These findings are in line with those of previous studies in which dietary stinging nettle at levels of 1-2% improved weight gain and FCR in broilers at rearing period (Safamehr et al., 2012). At the end of the experiment (day 35), the highest ADG belonged to the groups fed by supplemented diet with stinging nettle 2% and 4% compared to the HS and NC groups on the basal diet. Terpenoid phenols in stinging nettle protect intestinal epithelial cells from pro-apoptotic oxidant stress and increase epithelial cell growth, which improves the growth performance in birds under stress conditions (Gül et al., 2012). Additionally, the positive effects of the dietary stinging nettle powder on growth gain and FCR in the broilers exposed to oxidative stress was reported by Ahmadipour and Khajali (2019). The obtained results from the current study indicated that stinging nettle improved weight gain and FCR in the birds exposed to heat stress, which is important and economically valuable.

During heat exposure, birds are unable to regulate body temperature due to lack of sweat glands and abundant of feathers on their body. In response to high environment temperature, the body temperature of birds rises, and panting is a major physiological mechanism to dissipate the excess body heat by evaporating water from the respiratory tract. Bird panting is defined as an increase in the speed and range of respiratory movements (McKechnie et al., 2016). According to Table 6, the rectal temperature and panting frequency increased significantly in the heat-stressed groups compared to the NC. In agreement with these findings, Mehaisen et al. (2017) showed that heat stress increased rectal temperature and panting frequency in broilers. Based on the obtained results herein, the rectal temperature decreased in the broilers exposed to heat stress on the last day of heat exposure (day 29) compared to days 22 and 25, indicating the higher sensitivity of broilers to acute heat stress and the short time of chronic heat stress compared to long time. Our findings confirmed the results reported in a study by Rodrigues and Beldade (2020). They stated that broilers could regulate their body temperature in chronic heat stress; accordingly, the body temperature reduced in the last days of chronic heat stress compared to the first days. Stinging nettle supplementation at level 4% had a positive effect on the improvement of panting frequency and the rectal temperature on day 22 in the HS-SN4 group (first day of heat stress); however, it was not statistically significant (*P*>0.05). Even though the physiological parameters in this study were not significantly influenced by stinging nettle supplementation, the positive effects of antioxidants in regulating the physiological responses of the heat-stressed birds have been already proven (Mehaisen et al., 2017).

## Conclusion

Heat stress leads to oxidative stress and reduces growth performance in broiler chickens. The current study indicated that adding stinging nettle powder to diet improved the oxidative status of broilers exposed to chronic heat stress by decreasing MDA and HSP70, as well as increasing the T-AOC and activities of antioxidant enzymes. Furthermore, we revealed that the effects of stinging nettle were dose-dependent and supplementation of stinging nettle at level of 4% had stronger effects on mitigating the oxidative stress. In addition, this plant improved the growth performance in the heat-stressed broilers. Accordingly, it seems that dietary stinging nettle has a strong antioxidant and growth-promoting activities against heat challenge in broiler production industry. In addition, stinging nettle could be suggested to be utilized as a feed additive in the poultry diet for improving health status and defense mechanisms of birds under stressful conditions.

## Acknowledgments

The authors would like to appreciate Shiraz University, Iran, for their financial support (grant number: 98INB1M340866.)

## Conflicts of Interest

The authors declare no conflict of interest.

## Notes

### Competing Interest Statement

The authors have declared no competing interest.

